# Does evolution design robust food webs ?

**DOI:** 10.1101/663203

**Authors:** B. Girardot, M. Gauduchon, F. Ménard, JC. Poggiale

## Abstract

Theoretical works that use a dynamical approach to study the ability of ecological communities to resist perturbations are largely based on randomly generated ecosystem structures. In contrast, we propose here to asses the robustness of food webs drawn from ecological and evolutionary processes with the use of community evolution models. In a first part, with the use of Adaptive Dynamics theoretical framework, we generate a variety of diversified food webs by solely sampling different richness levels of the environment as a control parameter, and obtain networks that satisfactory compare with empirical data. This allows us to highlight the complex, structuring role of the environmental richness during the evolutionary emergence of food webs. In a second part, we study the short-term ecological responses of food webs to swift changes in their customary environmental richness condition. We reveal a strong link between the environmental conditions that attended food webs evolutionary constructions and their robustness to environmental perturbations. When focusing on emergent properties of our evolved food webs, especially connectance, we highlight results that seem to contradict the current paradigm. Among these food webs, the most connected appear to be the less robust to sudden depletion of the environmental richness that constituted their evolutionary environment. Otherwise, we appraise the “adaptation” of food webs, by examining how they perform after being suddently immersed in an environment of modified richness level, in comparison with a trophic network that experienced this latter environmental condition all along its evolution.

## Introduction

Understanding and linking both the structure and functioning of ecosystems has been addressed as one of the major issues in ecology, particularly due to increasing pressures on biodiversity (1, 2). The large diversity/complexity-stability debate (see (3, 4) for reviews), rooted in this issue, has been approached from different angles. A first angle tackled this complexity by studying food webs — the representation of who eat whom in ecosystems — since they provide tractable depictions of biodiversity and ecosystem structure and function (5). Pioneer food web models such as the cascade model (6) or the niche model (7) were able to produce satisfactory food web structures, with the ability to transpose some network properties from natural food webs. However, among the limitations of these models is first the fact that they do not propose an explanation of how these properties emerge, and second that they do not provide a quantification of interactions strengths, prerequisite for a population dynamics model of interacting species.

Past decade has seen the development of community evolution models, that came to adress these limitations (8–10). They provide demographic and evolutionary dynamics, presenting resulting food webs as the “winnowed survivors of evolutionary processes” (1). In those models, species are characterized by some traits (unique to multiple, depending on studies), which drive the demography of the whole interacting community on a given time scale. On another time scale, these traits evolve according to the selection pressures exerted on each of them. New species are introduced randomly, ultimately leading to the emergence of communities. The diversity and structure of the resulting food webs are hence emergent properties controlled by small scale species interaction parameters. The model developed by Loeuille and Loreau (8) has the advantage to be relatively simple as it implies only one characterizing trait for a species, the body size at maturity. However, networks that emerge from this model have been shown to underlie a very uniform regular structure, in particular because species have the same feeding characteristics (same feeding distance and same specialization level). In order to reproduce a more realistic diversity in feeding strategies and hence in food webs structure, it is necessary to consider these feeding parameters as evolving traits. Several studies do have attempted to consider the further evolution of these traits (e.g. 11).

Still with the aim of explaining the structure and functioning of food webs, it was noted that processes that operate at global scale (e.g environmental productivity, climate) have rarely been shown to be determinant for the structure of communities, even if some exceptions exist (12). Yet, several hypotheses have been formulated to explain how community structure could be related to environmental attributes, such as productivity and/or ecosystem sizes (13). Among properties of community structure, food chain length, or connectance, has been found to vary along environmental gradients, in lakes (13), streams (14) or high latitude marine ecosystem in a more recent study (15). To our knowledge, no studies using community evolution models have focused on the effect of parameters influencing these global-scale processes on the emergent structures, either because it was not their main topic, or because they led only to expected results. This is the case, for example in the seminal study of Loeuille and Loreau (8), in which discussions on the effects of variations in a parameter affecting the productivity of the environment in which the emerging network evolves (*i*.*e* global scale parameter) are brief, because they result in monotonous trends: an increase of specific diversity and maximum trophic level with richness of the evolutionary environment. Among the subsequent, more detailed community evolution model, such as Allhof’s model (11), none has investigated so far the effects of these global scale parameters on the structure of resulting communities. Although they are still in their infancy, community evolution models show great potential, especially because they can bridge the gap between ecosystem and community ecology (16). Among the promising but still unexplored avenues in the exploitation of such models is the analysis of both short-term ecological responses and long-term evolutionary responses of communities to disturbances (16).

A last angle mentioned here to broach the diversity-stability debate is to focus on how to relate the structure of food webs to their fragility. For example, motivated by the observations of increasing species losses (17, 18), several studies have investigated the links between primary species losses on communities, the so called secondary extinctions, and the structures of food webs (5, 19–21).

Here, we try to establish the link between these different approaches to the diversity-stability debate. In the first part, we build up a model inspired from (8), but modified by including two additional adaptive traits in order to bring out diversified feeding behaviors. The evolution component of the model is also improved and take opportunity of the Adaptive Dynamics toolbox. We then interested ourselves in the effects of the variation of a single control parameter that determines the richness of the environment during evolution (global scale processes parameter) and its effects on emerging structures. The properties of the food webs obtained in this way are then confronted to a large set of empirical data (22). In the second and main part of the article, we study the responses on the ecological timescale of the evolved food webs to abrupt changes in the richness of their evolutionary environment. We quantify the effect of the level of environment richness that constituted the evolutionary environment of a food on its robustness when it is subsequently confronted to a different condition. We then assess and discuss the possible direct links between emerged structure properties of evolved networks and their robustness. Lastly, we investigate whether an evolved food web performs the best in its evolutionary condition, compared to other networks that have first evolved in different richness levels and abruptly undergone this former condition. It contributes to the debate of considering the analogy of a food webs as a super-organism, that would become the fittest to its environment.

## Model

### Trophic network

The model is derived from Loeuille and Loreau’s work (8) (LL in the following), with, as we shall see later on, some modifications both in the ecological hypotheses and in the methods we used. For species *i* in a community, we denote *x*_*i*_ its typical (average, maturity) body size. Its feeding kernel is Gaussian-shaped and depends on two parameters: the preferred body size difference *d*_*i*_ between its own body size and the size of the species it consumes, and *s*_*i*_ the width of the kernel, so that it optimally consumes species of body size *x*_*i*_ − *d*_*i*_, with *s*_*i*_ determining its degree of generalism as a consumer. This setting allows for the determination of the connected species in the trophic network but also provides a weight on the connections. Thus, interactions in the trophic network are not set *a priori* but emerge from the relative body sizes of every species pair in the community, combined with their feeding kernels. As in subsequent papers (e.g. 10), we removed some limitations present in the original model.

### Adaptive Dynamics

For each species, we let the body size *x*_*i*_ and also the feeding kernel parameters *d*_*i*_ and *s*_*i*_ be adaptive traits that undergo evolution. We here depart from LL in which only the body size evolves, following some other authors that modified the original model (9–11, 23). The evolution of traits is driven by the selection pressures, which depend in turn on the network structure. But since network structure is determined by species traits, it changes as well on the evolutionary time scale and in turn reshapes the selection pressures on species. This is a school case of coevolution (24).

Adaptive Dynamics theoretical framework (25–27) is particularly well suited to encompass such situations in which co-evolution of traits can only be understood from a precise description of the selection pressures and where it feeds back on them. Adaptive Dynamics provides us a very powerful tool that allows us to model efficiently the coevolution of adaptive traits through a set of ordinary differential equations for expectation of traits values: the *canonical equation*.

### Population dynamics

Adaptive Dynamics key ingredient is the supplying of the *invasion fitness* fonction that allows to assess the invasion success of a mutant, according to its new trait value and the state of the resident community in which it appears. Invasion fitness function is built up on the explicit population dynamics of interacting species that both allows to determine the steady state of populations in the community and the initial dynamics of the mutant lineage just after it appears.

LL provides such population dynamics model from which we started on. Model is based on ordinary differential equations for species abundances. Main modifications from LL study concern (i) the limitation of the feeding kernel to only smaller species that we removed, what allows loops and cannibalism (ii) the functional response that is Holling-type II in our model, (iii) a simplified equation for the basal resource, and (iv) the introduction of two different feeding conversion coefficient, for the consumption of the basal resource or for the feeding on species.

For convenience, we denote 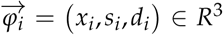 the trait vector, thus the phenotype of species *i*. Following this notation, the feeding kernel of species *i* is denoted 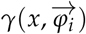 where *x* is the body size of the prey.

*N*_*i*_(*t*) denotes the biomass of species *i* at time *t. N*_0_(*t*) is the biomass of the basal resource. ***N***(*t*) = *N*_0_(*t*), *N*_1_(*t*),…, *N* (*t*) is the biomass vector of the community. 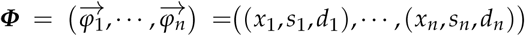 (*x*_1_, *s*_1_, *d*_1_),…, (*x*_*n*_, *s*_*n*_, *d*_*n*_) is the trait vector of the community. The population dynamics reads :

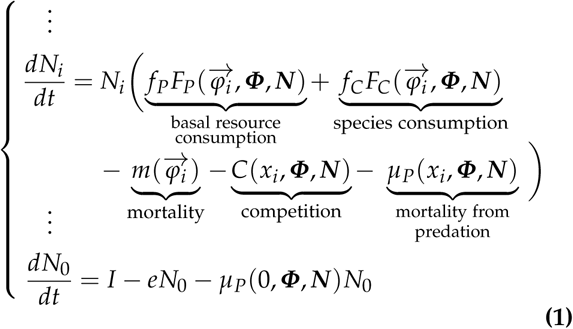

Table 1 gives the detailed formulations of the different terms of these differential equations.

**Table 1.**
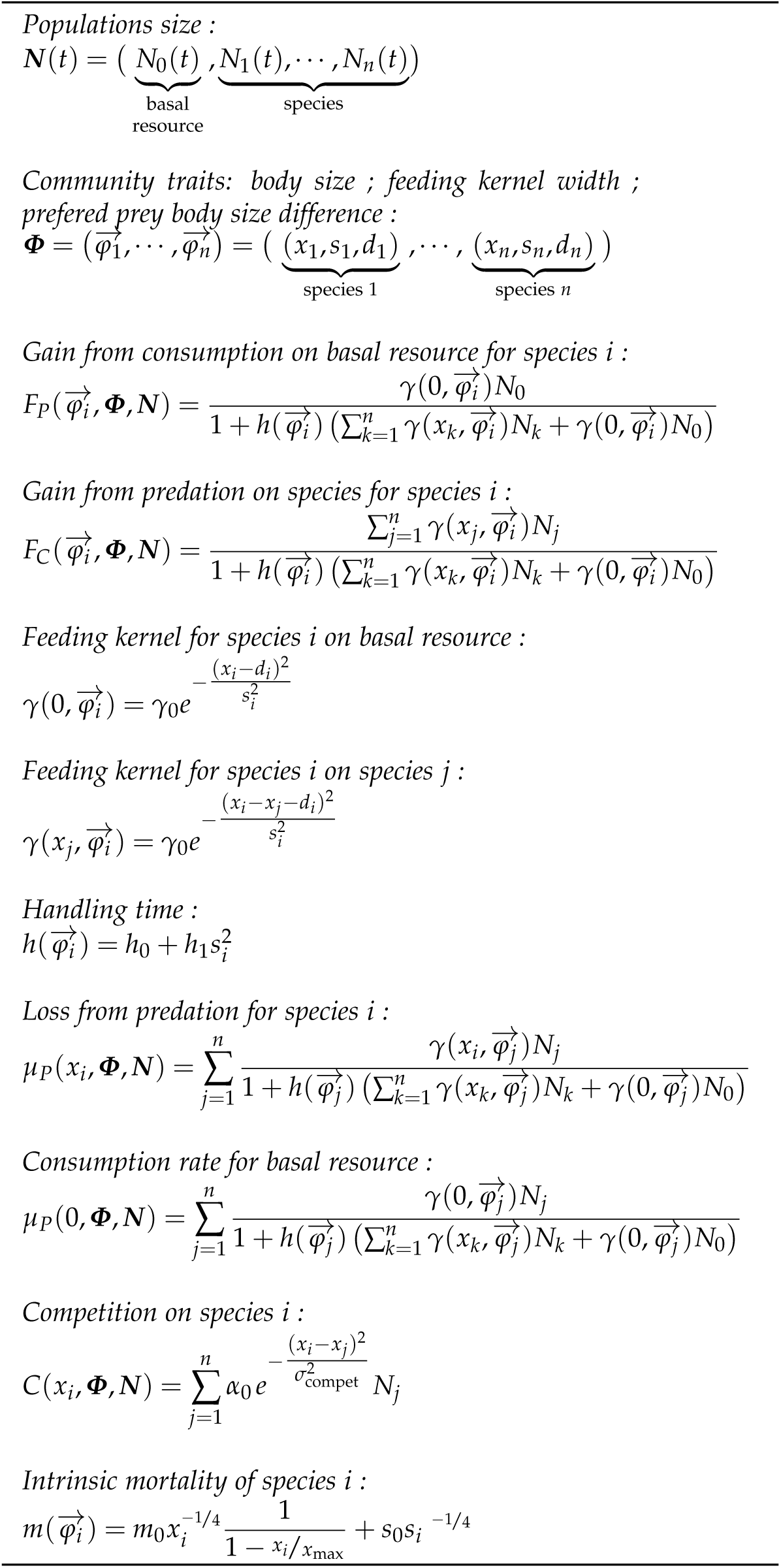
formulas

The choices in the parameter values are based on the existing literature. Parameters values for the resource dynamics are based on LL model, as well as competition and mortality functions parameters. As in (10) we replaced the body size dependent efficiency function by the two fixed assimilation efficiency parameters, one for basal resource, *f*_*C*_, and one for species resource, *f*_*P*_, following (28). Parameters values are indicated in the caption of Figure 2.

For the range of parameters values we considered, the joint dynamics of population sizes generally stabilizes, after a short transient dynamics, to a stable steady state.

### Canonical equations

The invasion fitness can be inferred from Eq. (1) :

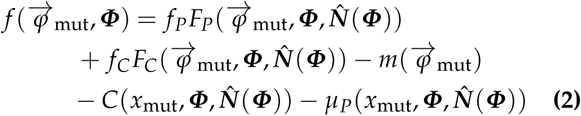

where 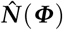 is the vector of equilibrium values of the populations biomasses, determined by the traits vector ***Φ***.

From the invasion fitness, Adaptive Dynamics toolbox let us infer the canonical equation for each coevolving species and for each trait *x, s* and *d*. For species *i* :

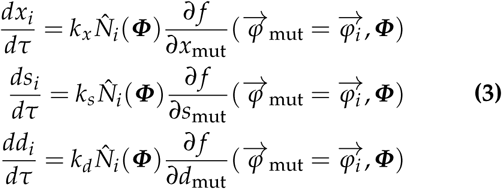

where *τ* is the evolution time and *k*_*x*_, *k*_*s*_ and *k*_*d*_ are mutation-related constants. This system of 3*n* ODE drives the coevolution of the *n* interacting species in the community. Evolutionary dynamics usually stabilizes to an equilibrium of traits values, an *evolutionary singularity*. But evolution does not necessarily stops here. One of the power of Adaptive Dynamics is that it can predicts, again in the basis of the invasion fitness function, the two subsequent alternative outcomes : long term stabilization of traits value, or the occurrence of the *evolutionary branching* that leads to the emergence of two new species in the place of their unique parent species.

### Trophic network emergence

We proceed with an initial single species (*n* = 1) for which we derive and integrate the canonical equations Eq. (3) for its three traits. If the evolutionary dynamics stabilizes to an evolutionary singularity that allows evolutionary branching, we replace the former species with two new species with slightly different traits compared to their parent species. We rewrite Eq. (3) with now 2 species (6 equations) and integrate it again.

We stop integration upon two possible events. (i) one of the species undergoes evolutionary branching: we replace it by two new species (ii) traits in the population co-evolve to a point where one of the population vanishes at equilibrium: we remove the corresponding species from the community. We rewrite Eq. (3) with the new number of species (*n* + 1 or *n* − 1).

We iterate the process until a chosen final evolutionary time is reached. A typical simulation is given by Figure 1 (panel II).

**Fig. 1.**
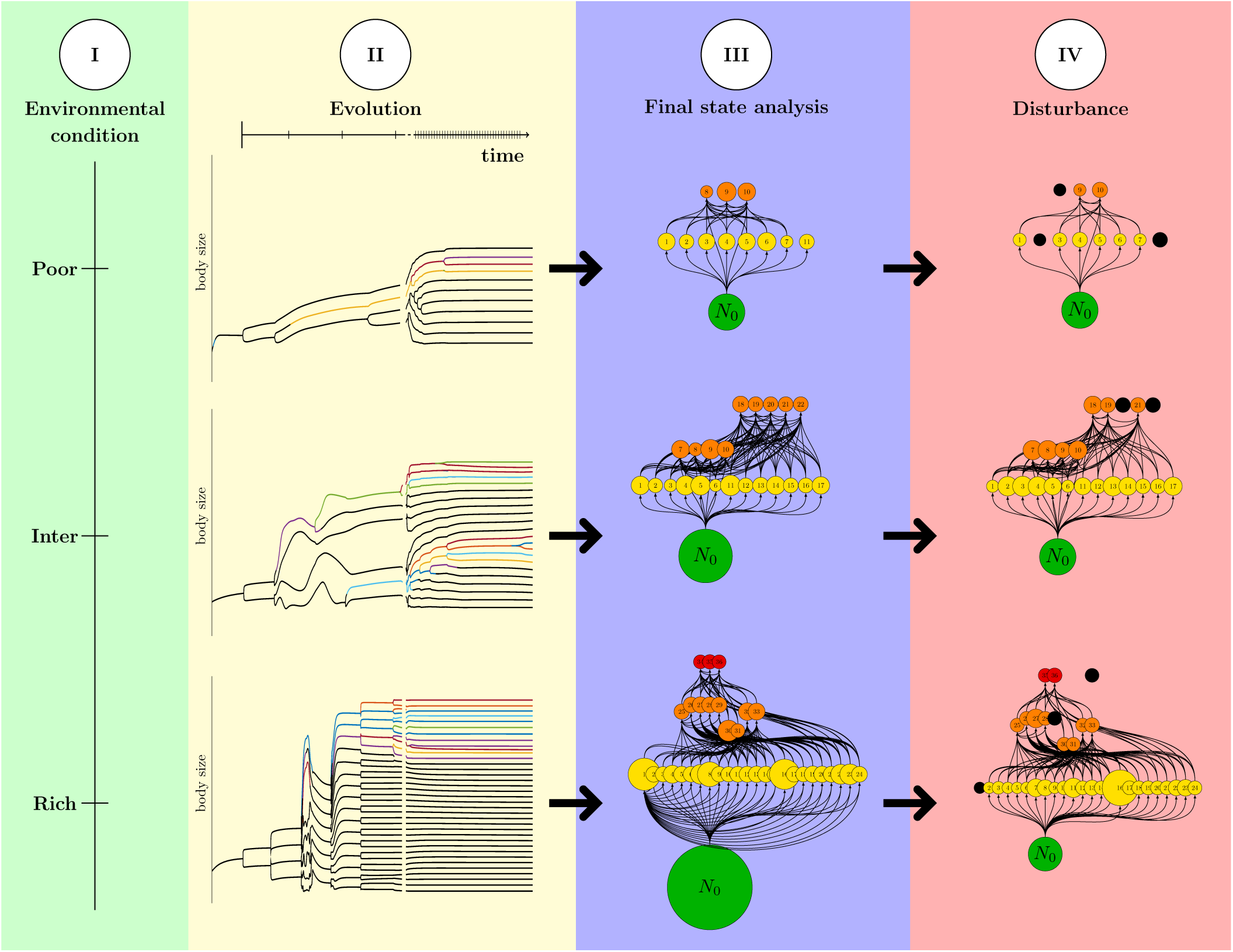
Explanatory scheme of the disturbance experiment. The first step consists in setting up the model for different environmental richness values *I* (panel I). We show here three example values : poor, intermediate and rich environmental conditions. The second stage (panel II) consists in simulating the eco-evolutionary model under these environmental conditions. Trophic networks emerge from succesive diversifications. Evolved food webs are analysed (panel III) for different emerging properties. Then (panel IV), they are confronted to changes in environmental richness, what impact populations on a short time scale, potentially inducing species extinctions (black circles).

This iterative approach of Adaptive Dynamics canonical equation has, to our knowledge, only been used but in a very recent study (29). Compared to LL, this simulation method is much more efficient: simulations run faster and by construction, species are well separated from each other so that the final states of simulation can easily been analyzed.

As already put out in (10), coevolution of the feeding kernel along with the body size allows the emergence of much more diverse trophic networks compared to models in which only body size evolves. On the other hand, as a downside, this additionnal freedom we inserted in the model released the individual parameters values (feeding kernel parameters *d* and *s* but also body size *x*) from the intermediate range they were confined into in LL’s model, so that we could not, as they did, neglect some of the constraints unveiled when parameters attain extreme values. We hence incorporated in the model additionnal costs due to high body size (high *x*) and to extreme specialization (very low *s*). Moreover, we modeled the handling time in the Holling-type II functional response as an increasing function of *s* so that a complete generalist consumers has a broader prey niche, but is less efficient in feeding than specialists. See table 1 for a precise formulation of these costs.

## Analysis of evolving food webs in a spectrum of contrasted environmental conditions

### Method

Trophic networks generated by evolution are highly dependent on the model parameters. Some of them are related to individuals or to interactions between individuals. Others are directly related to environment through the dynamics of the basal resource. We decided to focus our analysis on the global scale parameter representing the environmental richness, *I* (influx level of the basal resource, Eq. 1). This choice was first motivated by the fact that it allows to a certain extent to reproduce the contrasting environmental conditions of natural ecosystems. In addition, preliminary study shown that this parameter by itself allowed to generate a great diversity in simulated evolved food webs.

We explored some different values for the other parameters but then we fixed them.

For each value of *I* in a log scaled range of 1000 values from 0.1 to 350, we started from a single species, which traits (*x*_1_, *s*_1_, *d*_1_) were randomly chosen, and we let our algorithm generate an evolved, diversified, food web. We let the simulations run for 4.10^6^ time steps, what always allows the network to stabilize to some steady evolutionary final state.

Then we analyzed each evolved trophic network for a series of descriptors:

- species diversity *S* (final number of species),
- connectance *L*/*S*(*S*+1), where *L* is the number of realized links,
- maximum trophic level *TL*_max_, computed as in 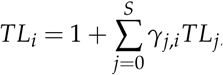,
- functional diversity indices for species attributes (body size *x*, generalism *s*, preferred prey body size *x* − *d*) ; simplifying formula from (31), we computed these indices as the weighted variance of the attribute:

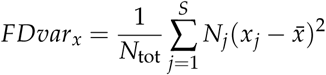

Figure 1 (panels I, II and III) synthetizes our method.

### Results and Discussion

We compiled data from 213 natural food webs from the ECOWEB database (22), and determine the properties of these food webs with the purpose to compare them to the properties of our theoretical emergent food webs. Figure 2 (A-D) confronts our theoretical networks to these natural food webs for the relationships between their specific diversity and their connectance, maximum trophic level, percentage of “top” species (consumed by no other species) or percentage of “bottom” species (that feed only on basal resource). The general trends we found were similar. Besides, although we only varied a single control parameter, the relationships between these emerging properties of evolved food webs are not curvilinear, the dots clouds really spreading over a 2D region. Even if natural data usually better fill the range of variation of some properties, we were able to reproduce a significant part of the diversity of trophic structures observed in the wild. This could suggest that the variability observed in the natural trophic networks might be explained by a limited number of factors.

**Fig. 2.**
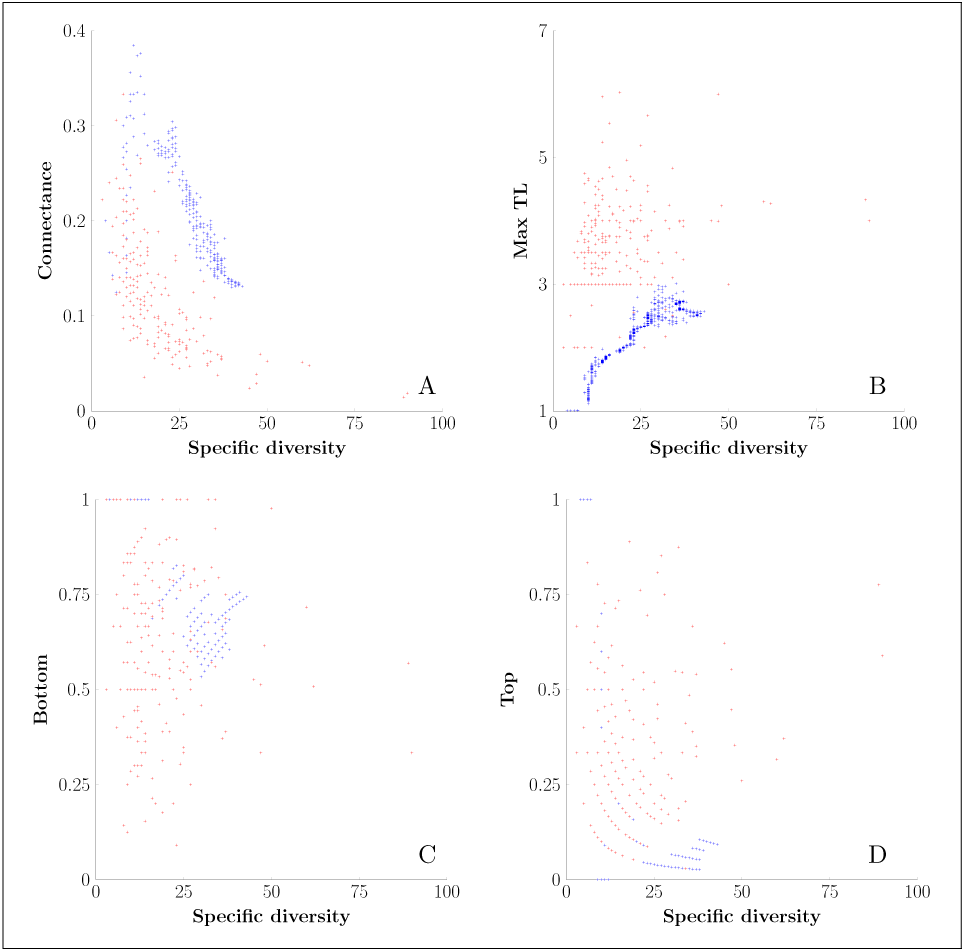
Emergent properties of evolved food webs (blue) compared to those of empirical food webs (red). Bottom corresponds to the percentage of species feeding on the basal resource. Top corresponds to the percentage of species that are not consumed. Parameters, expect *I*, are fixed and take the following values : *α*_0_ = 0.5, *σ*_*compet*_ = 0.4, *h*_0_ = 0.5, *h*_1_ = 0.05, *γ*_0_ = 1, *m*_0_ = 0.1, *x*_*max*_ = 15, *s*_0_ = 0.1, *f*_*p*_ = 0.85, *f*_*c*_ = 0.45.

We have not done more precise statistical tests here because it does not make much sense for several reasons. First of all, mere data fitting can be a poor test of model performance in many cases in ecology (32, 33) In addition, our species and their traits are not readily reliable with this set of empirical data as they do not consider body size and often pool several species in one node. Moreover, in empirical food webs, data are all the same compiled over a wide range of environmental situations, making it difficult to directly compare the variations of characterizing properties with a single environmental determinant as the environmental richness we focused on. Empirical studies that examine the links between such large-scale parameters and community structures do exist, though (15).

Figure 3 (A-C) shows in more details the variations in selected properties of emerging food webs against the gradient of environmental richness we considered. We chose to emphasize functional diversity indices for the adapting traits of species in the networks. The first striking result is that these indices of functional diversity, depending on which trait we measured them, shape differently. Functional diversity index on maturity body size *x* increases all along with environmental richness. Emerging specific diversity also follows the same trend, that is a monotonous increase with environmental richness (result not shown). Functional diversity index on generalism level *s* and on prefered prey body size *x* − *d* as for them are both maxized for intermediate values of the control parameter *I*. But the values of *I* that maximizes the functionnal diversity of *s* are low to intermediate whereas those maximizing the functionnal diversity for *x* − *d* are distinctly higher. Again, even if they do not appear on the figure, we additionnally tested connectance, which curves follows the shape of the functionnal diversity of *s* whereas the maximum torphic level follows the shape of the functionnal diversity of *x* − *d*.

**Fig. 3.**
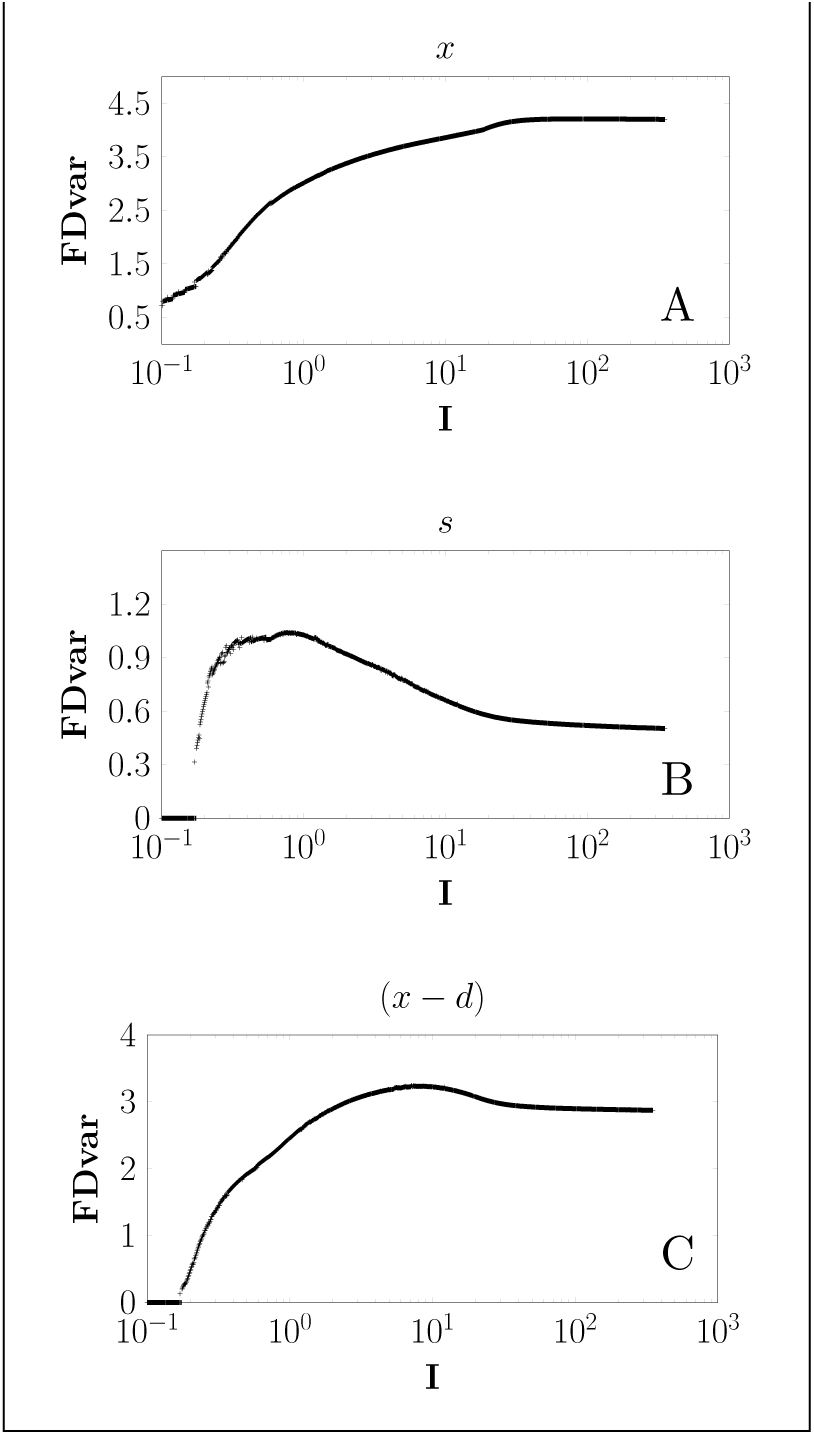
Functional diversity indices against the evolutionary environmental richness *I*. Functional diversity is computed for (A) body size, (B) degree of generalism, and (C) optimal prey body size.

The interesting point here is that these results make it possible to formulate hypotheses as to the key role of our control parameter, and only it, in the resulting emergent structure of food webs. In particular, three highlighted properties can not be maximized at the same time, because they are maximized when trophic networks evolved in three different environmental richness values. Highest specific diversity does not necessarily implies highest trophic complexities. This last point was previously highlighted in (30). Besides, previous studies have also investigated the role of environmental attributes on food web structure, and especially on food chain length. Three hypothesis have been formulated (13), namely the productivity hypothesis, the ecosystem size hypothesis, and the combined productive-space hypothesis. While this pioneering study shown a strong inclination for the ecosystem size hypothesis, other studies have shown results in favor of the other two. Our result on the complex structuring role of environmental productivity can give support to this productivity hypothesis. An advantage of our study is that we could isolate the effects of this structuring factor and test variations on a wide range of theoretical values. To illustrate this, we can compare our results with a recent study that looks at structural variations in marine communities at high latitudes along an environmental productivity gradient (15). Like them, we find the lowest connectance values for low environmental productivity, and higher connectance values, or higher trophic levels for high productivity values. However, in our case, the trophic complexity (connectance or functionnal diversity) of evolved food webs, after a first increase to some maximum value then begins to decrease again for high environmental productivity. This underline the fact that relationship between global scale parameters and network structure are not always straightforward and monotonuous and could give cues of why whichever simple hypothesis may be verified in some situations but defaulted in others.

## Disturbance experiment

### Method

There are numerous ways to understand and quantify the stability of an ecosystem (32). A large part of them consist in quantifying the resistance to perturbations. In this disturbance experiment, we exposed evolved food webs to new environmental conditions, different from the acustomary level of environmental richness during their evolution (Figure 1, all panels).

Effects of perturbations are assessed on a short time scale : once a food web has reached its evolutionary steady state, we abruptly change parameter *I* in the demographic system (1). It may only result in a change in the population abondances that stabilize, on the demographic timescale, to some modified equilibrium values. But it often happens that in this new equilibrium, one or several of the populations vanish, issueing species losses in the trophic network. We can therefore estimate how evolved food webs respond for various, new conditions, in terms of species diversity change (assessing species loss). But all other properties of food webs are likely to be impacted as well. We chose to focus on two properties. The maximum trophic level property, as in the previous section, and a new “information characteristic”: entropy. Entropy, or “information diversity”, is derived from Shannon’s information index (34), that quantifies here the diversity of trophic exchanges in a network (35–37). It is computed as:

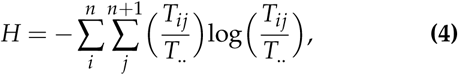

where *T*_*ij*_ denotes the magnitude of flow from node *j* to node *i*.

Each food webs, evolved in a richness level *I* = *I*_*evol*_, are confronted to a range of 1000 new richness levels *I*_*perturb*_ from 0.1 to 350. We hence get a 1000 × 1000 treatment. For each configuration, we determine the *relative species loss*, that is the number of perturbation-mediated extinctions, related to the pre-disturbance number of species. On contrasted situations, we assess the robustness of food webs, defined as the maximum intensity of disturbance they can undergo before they lose more than half of their species. We also calculate the gain in maximum trophic level, which can be positive or negative.

It is obviously difficult to determine *a posteriori* the environmental conditions an ecosystem has evolved in. This first approach allows us to determine how ecosystems react in face of changes in their environmental richness level, but we found the necessity to link an observable property of contemporary evolved food to their fragility or robustness. In a second approach, we chose to relate the connectance of evolved food webs to their relative species loss when confronted to a reduction factor in their richness level : *I*_*perturb*_/*I*_*evol*_ ranging from 0.2 to 1. As the connectance is an emergent property and not a control parameter, we used a smoothing gaussian kernel to merge data from irregularly distributed connectance values. The local variance of data we averaged with the smoothing kernel varies significantly and we quantified it precisely.

Lastly, we interested in the comparison of a trophic network that evolved in a given level of richness *I*_*evo*_ to other networks that evolved in other environmental conditions but were then subjected to this richness level as a perturbation. We compute the *comparative species loss*, corresponding to the difference between the number of species of a trophic network perturbed by a new richness level *I* and the number of species of a food web evolved from the beginning in this richness level *I*. Similarly, we computed the *comparative entropy loss*.

### Robustness of evolved food webs

Figure (4,A) shows the relative species loss for all 1000*x*1000 treatment configurations. It highlights three main outcomes. Overall, food webs appear to be quite robust to moderate environmental changes : no catastrophic collapses, relatively low relative species losses. Second, evolved food webs appear to be much more sensitive, in terms of species loss, to environmental depletion than to enrichment. Enrichment leads to a maximum of 10 − 15% species loss and often does not have any impact on specific diversity. Lastly, we found that food webs that have evolved in poor environments are less robust to environmental impoverishment than webs that have evolved in a rich environment. For a similar reduction factor in the environmental richness, food webs evolved in a rich milieu, even in terms of *relative* species loss, suffer much less of the perturbation than food webs evolved in poor conditions. The red arrows show the disturbances necessary to reduce the number of species by half, for two contrasted *I*_*evo*_. It happens for a factor 10 of reduction of then richness level for the left arrow whereas it needs a factor 200 for the right arrow.

**Fig. 4.**
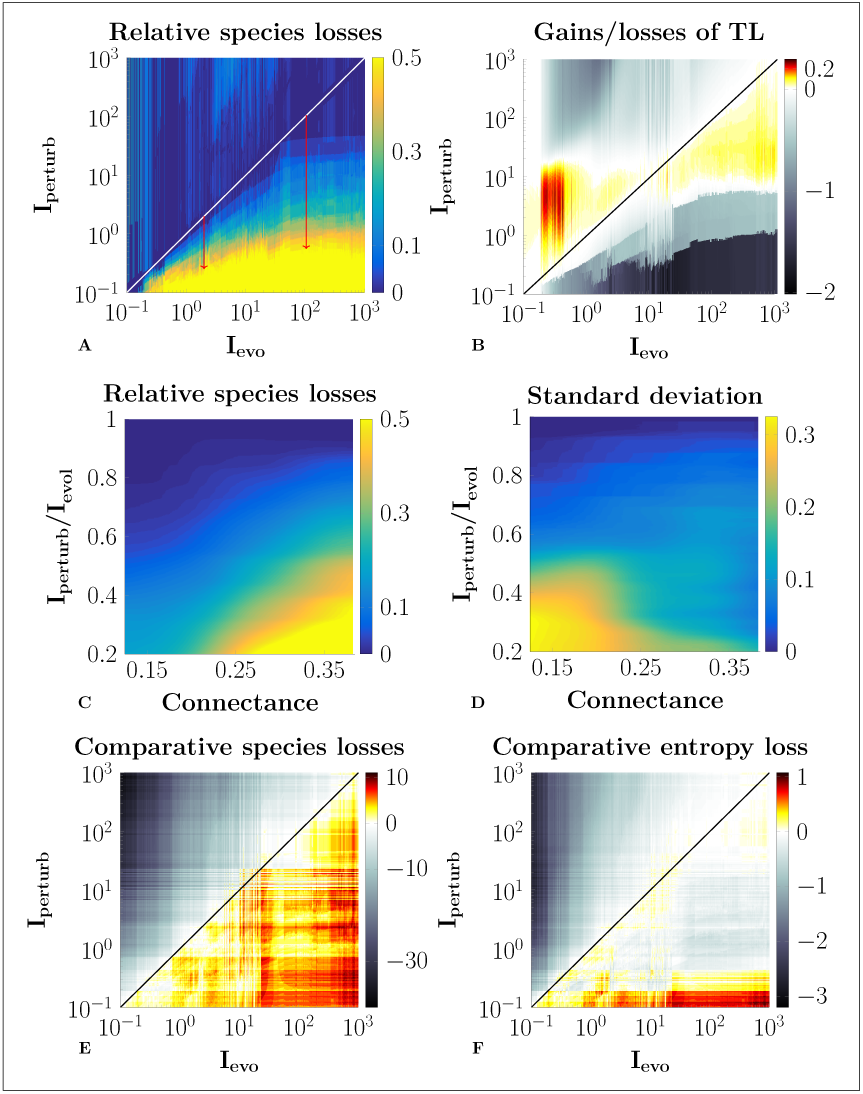
Short-term, ecological responses of food webs when they are subjected to new environmental conditions different from their evolutionary environment. (A,B) present respectively the relative species losses and the gains/losses of maximum trophic level observed. (C) shows the relative species losses when the productivity of the basal food web resource is depleted by a certain percentage. Food webs are sorted by connectance, and values are smoothed by a Gaussian kernel. (D) shows the local standard deviations of relative species losses. (E,F) show, respectively, the comparative losses of specific diversity and entropy (diversity of flows) of evolved trophic networks. Comparative loss refers to the difference between the new property value of a food web for a given, new, environmental richness, and the value of the same property for the “native” food web that has evolved in this later environmental richness.

When considering the maximum trophic level (Fig. 4,B), it appears, contrary to the results on species loss, that the evolutionary history parameter *I*_*evo*_ is not so significant any more, but that the maximum trophic level always tends to be maximized for some intermediate value. Indeed, trophic networks evolved in poor conditions increase their maximum trophic level when they are immersed in richer conditions, whereas for those evolved in rich environment, it increase when environment is impoverished.

Now relating the emerged connectance of evolved food webs back to relative species loss (Fig. 4, C), we focused on environment depletion. We observed that evolved trophic networks characterized by low connectance values appear to be more robust to an impoverishment of their environmental condition. Even for a division by 5 of the environmental richnes, they only lose 20% of their species. By contrast, the most connected of our evolved food webs lose about half of their species for only a division by two of the environmental richness. Yet, we assessed that the variance of the averaged species loss becomes important, espcially for low connected food webs experiencing strong perturbations (bottom left of Fig. 4, D). This results from the fact that a variety of evolved food webs, that experienced very different *I*_*evo*_ during their evolution, end up with convergent connectance values. Even if these low connectance food webs are on average much robust to perturbations, some of them, from our panel, evolved in poor environmental conditions and are highly impacted by further impoverishment.

Both topological and dynamic approaches were used to study community responses to species loss (5, 20, 21). Dynamic approaches were used, taking into account some of the weaknesses of topological approaches (e.g. misleading picture of potential secondary extinctions).

However, most of these are based on generalized LV models, ignoring the evolutionary history of the communities generated. Explicitely taking into account this evolutionary history, we get in our study a data set arguably closer to field observations.

The second type of divergence concerns the type of disturbance. These studies were carried out through the analysis of secondary extinctions following the initial loss of one or more species. Primary species losses are discussed as being the result of several disturbance factors, that however usually stay out of the scope of the studies. Even if the results may vary depending on how the species are primarily removed (e.g. primary or secondary consumers), one of the major findings of these studies is that food webs characterized by high connectance values appear to be the most robust.

Here we have a different approach. When primary extinctions are proximal causes for secondary extinctions, explicitly taking into account changes in environmental conditions is like considering distal causes for these extinctions. And hence, we do not decide *a priori* which species potentially disappear but we estimate it *a posteriori* from the new environmental parameter value along with the population dynamics model.

And therefore, our apparently opposite results, because they were obtained on a selected data set and with a different methodology, actually do not contradict the current paradigm around the connectance-robustness debate. Instead, we provide additional elements of understanding and underline the precaution of attributing a high robustness to highly connected food webs if the environment is to undergo depletion disturbances. We also highlight that food webs with similar level of connectance may respond very differently to perturbations so that the average response may not always be very informative. This final result is obviously very sensible on the data set used and we recommend that precautions be commonly taken.

Another advantage of the framework used here is that it allows us to look at the impacts of an explicit disturbance, here the change in environmental richness, on other aspects of biodiversity than specific diversity. In particular, studying trophic level loss/gain can be allowed because we have access to the demography of the different species, which is not the case with topological approaches for example.

In our setting, perturbations do not change trophic connections between species. The results where maximum trophic level increases due to perturbations could only be explained because demography was rebalanced. For example, for cases where maximum trophic level increases with richness reduction, biomass of low level species, that exclusively consumes basal resource, decreases, thus representing a reduced proportion of the diet of higher level consumers, whose trophic level therefore increases.

### Adaptation of evolved food webs

Results on maximum trophic level (Fig. 4,B) shows that evolutionary history do not always shape food webs in a way that their functionning, at least for some characteristics, be optimized for the environmental conditions that have accompanied its evolutionary emergence. Indeed, all food webs increase their maximum trophic level when they are immersed in some optimal intermediate level of richness *I*_*opt*_, regardless of their evolutionary conditions.

Undoubtedly, a food web is not an adaptive unit that directly undergoes natural selection. Still, we wanted to push this analogy and assess whether trophic networks could be designed by co-evolution so that they would particularly fit their environment for some other property. With that goal, we observed our setting as a *common garden* experiment: we compared food webs “transplanted” in a given environment, one of them evolved in the same conditions, the second one in an other condition, richer or poorer.

Figure 4 (E) shows the comparative species loss of the “exogenous” food web compared to the “native” food web. The striking result is that, concerning specific diversity, food webs that emerged in poor or intermediate level of richness are not especially best “adapted” to their environment. Indeed, most of the trophic networks that evolved in a richer milieu, and then gained a greater specific diversity, keep a higher number of species when they undergo even severe environment impoverishment, compared to food webs evolved from the beginning in these poor environments (red area under the diagonal).

Figures (4,E,F) allows us to estimate whether the evolutionary history of a food web leads it to be more adapted to its evolutionary environment than any other food web that undergoes these conditions after its emergence. With regard to specific diversity, it appears that food webs that have evolved in rich environments and are then confronted to an impoverished environment have more species than food webs that have adapted to this poor environment during their evolution. It therefore appears that evolved trophic networks do not seem at first sight to be the most suitable for their evolving environment, at least not in view of the specific diversity present.

For connectance, not presented results show a symmetrical situation: networks that have evolved in poor environments, which then subsequently enriched, have (still) higher connectance than networks that have evolved in these rich environments. It appears that the conclusions of which is best fitted to a given environment thus may vary according to the properties studied.

However, when looking at properties derived from information theory, we found that evolution could optimize food webs for some characteristics. Indeed, the properties related to the organization and diversity of flows within evolved trophic networks show this trend. For most evolved food webs, any disruption of environmental richness from accustomary conditions induces values of these properties that are lower than those that characterize food webs that have evolved in these environments (Fig. 4, F). This means that the structure of food webs and the flows within them seem optimized by evolving processes.

The fact that ecosystems may develop or evolve by maximizing some of their properties is a long, contentious debate in ecology (32). The approach discussed here allows us to question the level of adaptation of a food web to its evolutionary environment by comparing it to other food webs subjected to this environment without an adaptation period. This provides a complementary point of view to the extent to which evolution leads an entity, here, in this case, a food web, to develop optimal properties. Considering food webs as super-organisms is a concept widely discussed in the evolutionary and ecological literature (32, 38). Loreau (32) argues that while near optimization of ecosystems properties is conceivable (with constraints on strengths and directions of individual and group selections), there is no reasons to believe that it will be generally achieved, and that there is no objective criterion to forecast it. Here we present contrasted results. Contrary to what usually show cross-transplantation experiments on real organisms, for food webs, some of them are better fitted to some environments than other food webs “adapted” to them (results on number of species, connectance). Besides, ecosystem properties related to diversity and organization of flows appear to be the best candidates for a generalized near-optimal behavior.

## Conclusions

In this paper, we first focus on the influence of a global scale parameter, environmental productivity, on the structure of food webs. Using a community evolution model, we look at the extent to which variations in this control parameter alone can lead to the emergence of food webs that reflect the diversity of structures observed in the wild. We show that a considerable part of this diversity can be reproduced in this way, underlining the importance of environmental productivity as a determinant of community structure. The second particularity of this study is to study the shortterm ecological responses of the food webs produced in this way. We focus on an explicit disturbance mode, which aims to modify the control parameter once the evolution step is completed, and thus quantify *a posteriori* the losses (or gains) of different network properties. Food webs characterized by high connectivity values appear to be the least robust to a decline in their environmental condition, given the resulting percentages of species loss. The induction of intermediate environmental productivity values positively (or neutrally) affects all food webs, with respect to the maximum trophic level encountered. This disturbance experiment finally allows us to question the potential status of food webs as “super-organisms”, i.e. their state of adaptation to their evolving environment, by comparing and contrasting other food webs subject to this environment. Some properties, linked to the diversity of flows within networks, appear to be the best candidates for possible properties that would thus be maximized by evolutionary processes. We believe that this work represents a first step in the unexplored avenue of exploiting community evolution models to analyze responses to disturbances of such winnowed survivors.

## Acknowledgments

We thank Clement Aldebert for helpful discussions and comments. This article is part of the CIGOEF French ANR. The PhD scholarship of BG was funded by the French Ministry for Education and Research.

## Authors contribution

B.G. and M.G. designed the model, performed the study and analysed the results. All authors contributed to the writing of the manuscript.

The authors declare no conflict of interest.

